# Signatures of Jamming in the Cellular Potts Model

**DOI:** 10.1101/2023.07.10.548321

**Authors:** Alexander J. Devanny, Daniel J. Lee, Lucas Kampman, Laura J. Kaufman

## Abstract

We explore the jamming transition in the Cellular Potts Model (CPM) as a function of confinement, cell adhesion, and cell shape. To accurately characterize jamming, we compare Potts simulations of unconfined single cells, cellular aggregates, and confluent monolayers as a function of cell adhesion energies and target cell shape. We consider metrics that may identify signatures of the jamming transition, including diffusion coefficients, anomalous diffusion exponents, cell shape, cell-cell rearrangements, and velocity correlations. We find that the onset of jamming coincides with an abrupt drop in cell mobility, rapid transition to sub-diffusive behavior, and cessation of rearrangements between neighboring cells that is unique to confluent monolayers. Velocity correlations reveal collective migration as a natural consequence of high energy barriers to neighbor rearrangements for certain cell types. Cell shapes across the jamming transition in the Potts model are found to be generally consistent with predictions of vertex-type simulations and trends from experiment. Finally, we demonstrate that changes in cell shape can fluidize cellular monolayers at cellular interaction energies where jamming otherwise occurs.

## Introduction

Coordinated cell migration is critical to a variety of biological functions including cell migration, morphogenesis, and cancer invasion^1, 2^. For example, during morphogenesis tissues experience dramatic changes in structure as cells rearrange, moving in collective swaths as complex tissues form and take shape^3, 4^. For these processes to occur, cells must be able to shift between relatively immobile and highly mobile states. The ability of cells to move in a coordinated fashion is necessary for the shaping and reshaping of tissue, while the ability to remain in place is essential to the long-term stability of a tissue. Complex signaling pathways underlie these cell motility programs^5^.

Broadly, a transition between high and low mobility states may be described as a jamming transition, a switch between a liquid-like and disordered solid-like state in which the dynamics of the system slow dramatically. Jamming was initially described for a system of frictionless, deformable spheres where a transition from free diffusion to collective motion preceded the abrupt arrest of the system components as a function of decreasing temperature, increasing density, or decreasing shear stress^6–8^. Active systems, including epithelial cells, have been shown to undergo analogous transitions from diffusive behavior to collective motility that precedes arrest to a solid-like state^9–11^. In recent years, the jamming transition has been extensively investigated in such active systems, with particular focus on understanding its implications in biological processes^12, 13^

In biological systems, the jamming transition has been understood to describe a precipitous dynamical slowdown and/or arrest of motion of the system components and has been investigated in a number of experimental and theoretical contexts. The transition has been shown to have clinical relevance in asthma, where primary cells from asthmatic donors were characterized by their resistance to jamming relative to non-asthmatic donor cells^14, 15^. Unjamming has been implicated in wound healing and cancer progression^16, 17^. Collective motility programs observed in wound healing are inherently connected with glassy behavior in cells^18^. In cancer, epithelial cells undergo the epithelial to mesenchymal transition (EMT) in which cell contacts are disrupted and cells become migratory^19, 20^. In 2D and 3D cell culture, cells that have progressed through EMT display fluid-like behavior while healthy epithelial cells remain stationary, as in a solid^21–24^. However, it remains unclear to what extent unjamming and EMT are distinct, and to what extent one or both are required in cancer invasion^20, 25, 26^.

For cellular systems, a hypothetical jamming phase diagram has been proposed, with jamming occurring as a function of increasing cell density, increasing cell-cell adhesion, and decreasing myosin dependent-motile forces^27^. Effective cell adhesion is proposed to be governed by an interplay of adhesion and cortical tension that is critical to determining the energy barriers to cell movement that dictate fluid or solid-like behavior^28, 29^. Here, cadherins primarily dictate the degree of cell-cell adhesion while actomyosin contractility dictates the strength of cortical tension opposing the extension of cell-cell contacts^30, 31^.

The vertex model, and extensions thereof, has proven to be a popular method and powerful tool for probing the jamming transition in biological systems^32–34^. In vertex model simulations, cells are composed of a series of vertices and edges, which evolve according to an energy functional. These simulations shed light on the forces at play in epithelial sheets that lead to the cellular dynamics observed during migratory processes as well as jamming. Vertex-type simulations have specifically highlighted the importance of cell shape in the jamming transition^35, 36^. In particular, it has been shown that the jamming transition occurs at a characteristic value of cell shape index, the dimensionless ratio of perimeter to area, P/A^1/2, 34^ and the distribution of cell shape indices correlates with liquid- or solid-like behavior of cells^37, 38^. Experimental work has further confirmed the value of cell shape index as a structural descriptor of jamming within a collection of cells – one that is both easily accessible and universal across biological systems^37^.

The Cellular Potts Model (CPM), a type of lattice-based Monte Carlo simulation, has been a staple of cell migration investigations since its introduction but has only rarely been used to investigate jamming^39–43^. The advantages of this model are its relative simplicity and access to sub-cellular detail. Despite its simplicity, the CPM has seen use in diverse contexts such as avascular tumor growth and cell migration, including as a tool to study both cell-matrix interactions and forces exerted during migration^44–48^. The CPM is an extension of the Q-Potts model, in which cells are modeled as an assemblage of contiguous like-spins on a lattice. Adjacent unlike-spins are assigned energies of adhesion, and the CPM stochastically evolves to minimize the energy of the system according to a Hamiltonian typically composed of area, perimeter, and adhesion energy terms. Cell motility arises from spin flips during each Monte Carlo step (MCS) that are accepted with a conditional probability associated with the change in total system energy. Most CPM implementations use similar forms of the Potts Hamiltonian, though variations are sometimes seen in which the area or perimeter term is not included^48, 49^ or in which additional terms are added to capture more complex cell behaviors^40, 50, 51^. CPM studies of cell migration have recapitulated aspects of experimental epithelial collective cell migration and, recently, the CPM was used to demonstrate glassy dynamics of cellular monolayers reminiscent of the jamming transition^42, 43^. Taken together, these works suggest that the CPM could also be a useful tool to investigate cellular jamming. Towards this end, it is necessary to perform a more rigorous analysis of the model in the context of indicators of jamming identified by and used in vertex-type models and experimental studies, as well as make connections between collective migration and jamming in Potts model studies.

In this work, we employ the CPM to investigate the jamming transition by analyzing the motility of cells under varying degrees of confinement, from unconfined cells, to aggregates, to confluent monolayers. We find cellular monolayers display a distinctive abrupt drop in diffusivity as a function of cell adhesion, with jamming marked by a transition to sub-diffusivity. Cell shape indices in the CPM are found to be similar to and generally consistent with predictions of jamming in other model and experimental systems. Within the CPM, we propose an approach for characterizing cellular rearrangements as a metric of jamming analogous to explicitly defined cell rearrangements in the vertex model. Velocity correlations in cell monolayers show increasing spatial correlations indicative of collective motion as cell adhesion energies increase, with a maximum in this collective motion directly preceding cell arrest. Finally, we demonstrate that increasing cell shape index in dense monolayers is sufficient to fluidize an otherwise jammed monolayer, and that a combination of mobile anisotropic cells and otherwise arrested rounder cells can fluidize a monolayer independently of cell-cell adhesion energy, demonstrating the potential impact of EMT-induced shape changes on cell motility.

## Model

### Cellular Potts Model

To investigate jamming, we utilized a two-dimensional CPM. It consists of *N* cells, existing on a square lattice of dimensions *L* x *L* (*L* = 100), with periodic boundaries, each of which has a unique spin integer, σ(*i*, *j*) ∈ {1, 2, …, *N*}, where (*i*, *j*) identifies the position of a lattice site. The environment is represented by the spin of σ(*i*, *j*) = 0. Sub-domains with identical spin σ ≠ 0 represent a distinct cell, and they can be divided into cells of a particular type τ(σ). The simulation runs by Monte Carlo methods, using a modified Metropolis algorithm^52^, with iterative sequences of spin flips that attempt to minimize the total energy of the system. Each time step, or Monte Carlo sweep (MCS), is composed of *L*^2^ flips. For each flip, the spin of a randomly selected lattice site σ(*i*, *j*) is changed to the spin of one of its neighboring sites σ(*i*′, *j*′) of different spin integer (σ(*i*, *j*) ≠ σ(*i*′, *j*′)). A flip is accepted with probability given by:

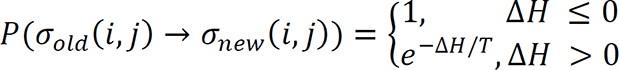

where *H* is the Hamiltonian of the system. The Boltzmann weighted probability function guides the simulation so that there is a stochastic reduction in the total energy of the system. The Hamiltonian at any time, *t*, can be divided into three components:

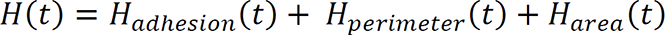

The adhesion component is based on the differential adhesion hypothesis^53^ and, in the simplest cases, may be related to number of active cell adhesion molecules per cell surface area. The adhesion energy is given by:

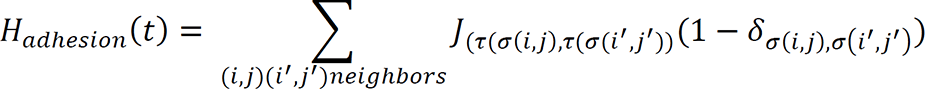

where J_(τ(σ))_ values are adhesion energies associated with cell-cell or cell-environment interactions, and higher J values are associated with a greater energy penalty associated with cell contacts. The perimeter and area components constrain cell shape and size, assigning a penalty for deviations from target values. These terms are given by:

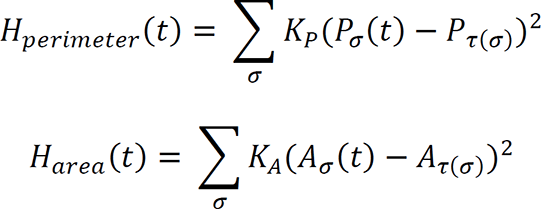

where *K*_P_ and *K*_A_ are constants determined empirically. *P*_σ_(*t*) and *A*_σ_(*t*) are the perimeter and area of a cell σ at time *t*, while *P*_τ(σ)_ and *A*_τ(σ)_ are the target perimeter and target area for each cell type, τ(σ), respectively. Target area and target perimeter reflect cell incompressibility and cortical elasticity, respectively, while the constant multipliers reflect the strength of the constraint. While these terms control cell shapes by constraining perimeter and area, *H*_adhesion_(*t*) can also influence cell shape. Parameters used in the simulations described within are provided in Table S1.

For simulation of isolated cells, individual cells were deposited randomly on an empty lattice. Simulations were run for 5 MCS to allow cells to equilibrate to reasonable shapes after deposition. For simulation of cell aggregates, cell centers were randomly deposited in a circular region in the center of the lattice. A Voronoi tessellation was generated from the centroids to divide the region into cells. The resulting tessellation was discretized to a 100 × 100 lattice for simulations. As in isolated cells, simulations were run for a short time (100 MCS, in this case) to allow cells to equilibrate to reasonable shapes after deposition. Equilibration times are short but sufficient to attain reasonable cell shapes for each simulation in the cases described above. After this initial equilibration, simulations were run for 1000 MCS or 100,000 MCS, for isolated cells and aggregates, respectively. For monolayer simulations, cell centers were randomly deposited across the full 100 × 100 lattice, and a Voronoi tessellation was used to construct cell shapes from these centroids, as in aggregates. Monolayers were similarly initially run for normalization of cell shape, here for a longer period of 1000 MCS due to the more crowded system, then run for a total of 100,000 MCS. Isolated cell simulations were run such that there were 20 separate individual cell trajectories for analysis. Aggregate and monolayer simulations were run in triplicate (150 and 300 total cells, respectively) for each parameter set.

## Results

### Cell Adhesion in Three Cellular Contexts

The cellular interaction energy, J, is a critical component of the CPM that has a substantial effect on cell behavior, as J encodes information on adhesion. Given the well-documented effects of adhesion in cellular jamming, this is a key component of interest when exploring the parameter space of the CPM. To characterize jamming within the CPM, we performed a parameter sweep of the interaction energy from J = 0 to J = 4, with higher interaction energies representing higher energy penalties for adding new cell contacts. Most experimental and theoretical studies of jamming consider the process in monolayers, with packing fraction equal to unity. To separate the effects of monolayer-induced confinement from the CPM-specific effects of varying J, we performed this parameter sweep in three different contexts: isolated cells, aggregates, and confluent monolayers (**Fig. 1**). Isolated cells, as the name implies, involves simulating cells that are sufficiently far from neighbors such that the cells can change shape and move through the surrounding environment without contact with another cell. Aggregates consist of a central cluster of cells surrounded by empty (environmental) lattice sites and approximate spheroid models that are often used to study cancer cell invasion^54, 55^. A confluent monolayer is a continuous layer of cells with no empty lattice sites. For simulations of both isolated cells and cell aggregates, environmental (i.e. cell-free) lattice sites are present; thus, over the three types of simulations, definition of two interaction energies is required – J_cell_, which dictates the energy penalty for cell-cell contacts, and J_env_, which dictates the energy penalty for cell-environment contacts.

**Figure 1.**
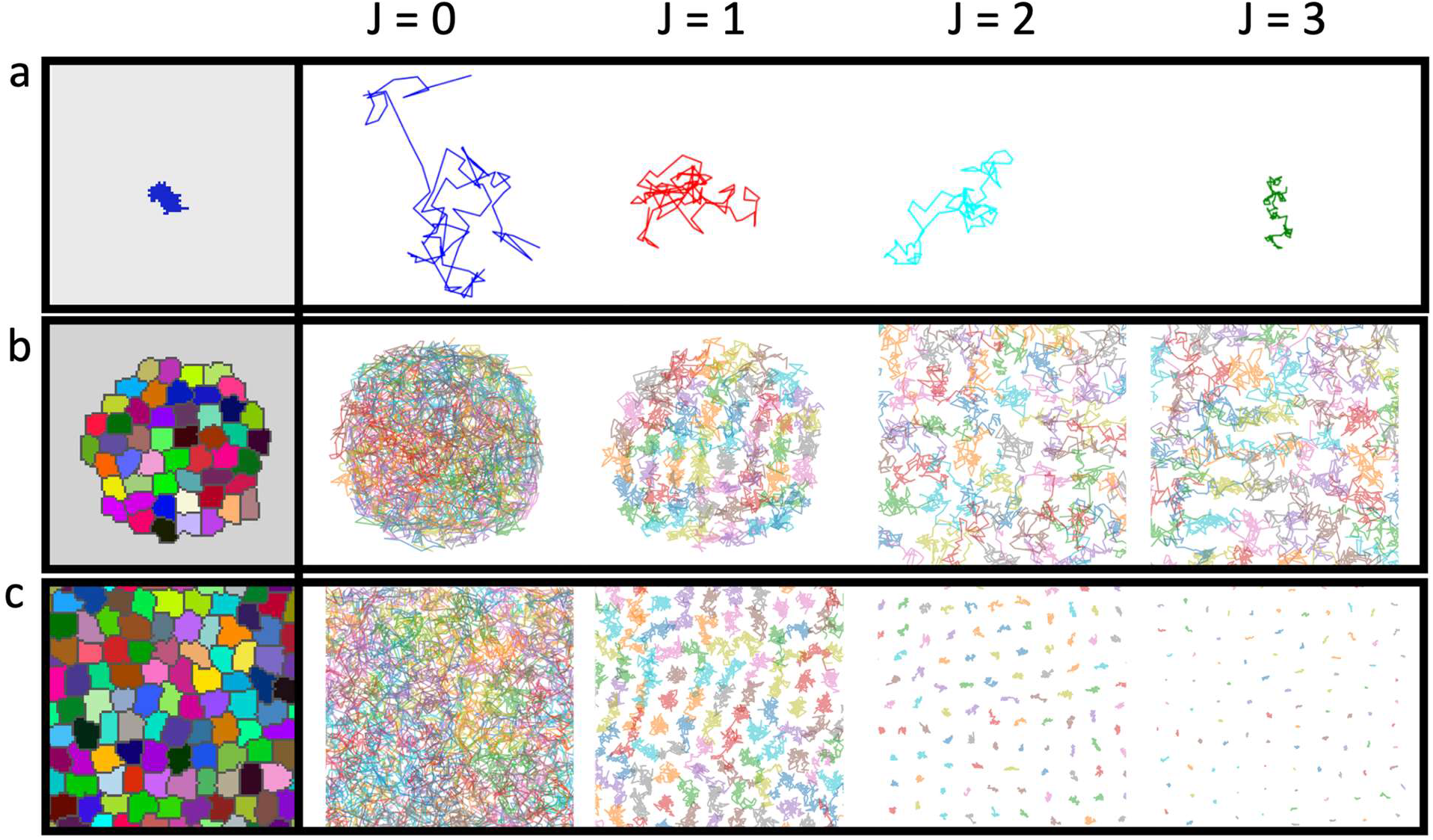
Potts simulations of single cells, aggregates and monolayers for a variety of J values. At left, exemplary cell positions for a (a) single cell, (b) aggregate, and (c) monolayer. At right, representative (a) single cell trajectories with (left to right) J_env_ = 0, 1, 2, and 3, (b) aggregate trajectories for J_cell_ = 0, 1, 2, 3, all with J_env_ =2, and (c) monolayers trajectories with J_cell_ = 0, 1, 2, 3. The trajectories shown for each case cover (a) 1000 MCS or (b-c) 10000 MCS. All trajectories are shown on the same size square lattice (100 × 100).

For isolated cells, J_env_ is the only adhesion energy that affects the system energy, as no cell-cell contacts are present. Varying J_env_ allows direct examination of the effect of interaction energy as a barrier to cell movement. Qualitative examination of cell trajectories reveals a steady decrease in cell motility as adhesion energy increases, with spin flip acceptances becoming less likely as dictated by the Boltzmann-weighted flip acceptance probability in the CPM (**Fig. 1a, Video 1**). In aggregates, the adhesion parameter that most strongly affects cell motility is J_cell_, as all cells initially have contact with neighbor cells. As such, we fixed J_env_ = 2 and varied J_cell_ from 0 - 4. When J_env_ is sufficiently high relative to J_cell_, cells are confined to a roughly circular aggregate as cells adopt more cell-cell contacts to minimize the unfavorable interactions with the environment. Qualitative examination of simulated cells in aggregates reveals extensive overlap of cell trajectories at the lowest J_cell_ = 0, followed by caging behavior at J_cell_ = 1 (**Fig. 1b, Video 1**). As J_cell_ approaches J_env_, caging behavior ceases, and fluctuations allow the eventual, slow disintegration of the aggregate (J_cell_ = 1.75, **Video 2**). When J_cell_ ≥ J_env_, disintegration is much more rapid, as cells repel one another and move into initially unfilled regions of the lattice (**Fig. 1b, Video 1**). For any multicellular situation in which cell-cell and cell-environment contacts are important, the relative magnitudes of these quantities will dictate the stability of the aggregate.

In confluent monolayers, the adhesion energy in the system is described by the single interaction energy J_cell_, as all lattice sites are filled with cells. For cells in the confluent monolayer, as in the case of aggregates, cell trajectories overlap significantly at low J_cell_, indicative of significant cell motion and exchanges of neighbors. Apparent caging is observed as J_cell_ is increased (**Fig. 1c, Video 1**). There is a modest onset of caging observed at J_cell_ = 1 in monolayers, with much stronger caging seen at values of J_cell_ = 2 and beyond.

### Cell Adhesion and Sub-Diffusion in the Potts Model

Cell trajectories were analyzed in greater detail by examining mean squared displacements (MSDs). MSD was calculated via 〈MSD〉 = 〈r(t) − r(0)〉, where brackets denote averaging over cells but not time lags, as such temporal averaging may mask sub-diffusion^56^. Without temporal averaging, ensemble averaged MSDs can suffer from noise. To improve the confidence of subsequent fits, we utilized the mean maximal excursion (MME) method, described in more detail in Ref. ^57^. MME curves report similar dynamics and give better quality fits than their MSD counterparts (**Fig. S1**). For extraction of the anomalous diffusion exponent, MME curves were fit to a power law of the form, MME = Kt^a^, with ⍺ = 1 indicative of normal diffusion, > 1 indicative of super-diffusion, and < 1 indicative of sub-diffusion. To determine diffusion coefficients, MSD was recalculated via 〈MSD〉_T_ = 〈r(t + τ) − r(t)〉_T_, where brackets denote averaging over cells and *T* denotes averaging over time lags τ. The early segments of time averaged MSDs were linear upon inspection; thus, we determined a diffusion coefficient D via a linear fit to the first 6 points of the MSD curves, with MSD = 4Dt. We characterized the diffusion coefficient and anomalous diffusion exponents across all contexts (isolated cells, aggregates, and monolayers) and varied J values. For isolated cells, the reduced cell motility seen in cell trajectories is reflected in the MSD curves and associated diffusion coefficients (**Fig. 2a,b**), as evidenced by an approximate order of magnitude decrease in the latter as J_env_ increases from 0 – 4. Despite this decrease, cells remain normally diffusive at all J_env_ values (⍺ ≈ 1, **Fig. 2c**).

**Figure 2.**
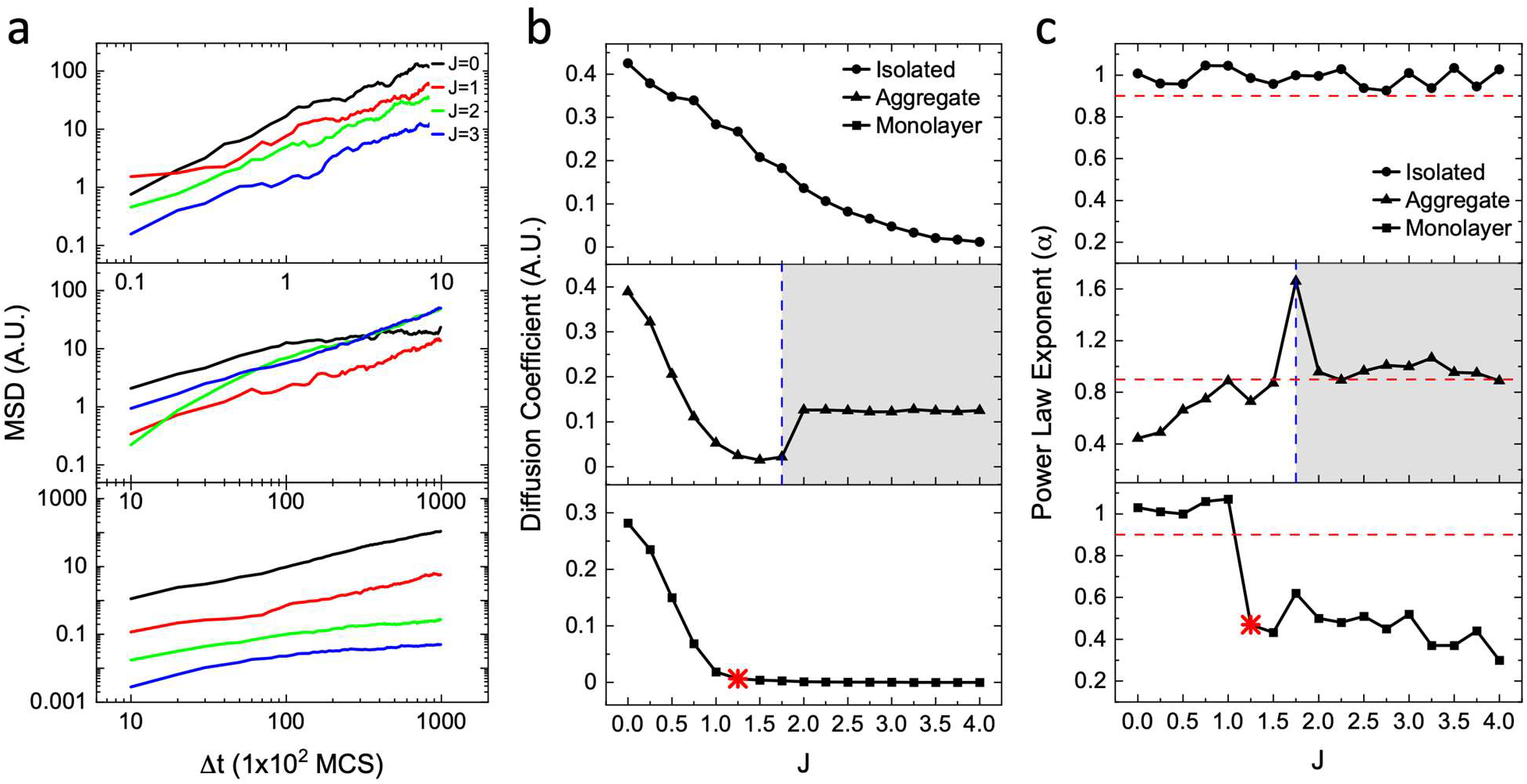
MSD analysis of Potts model simulations as a function of the adhesion parameter J. For isolated cells, J refers to J_env_, while for aggregates and monolayers this refers to J_cell_. (a) MSD plots for select J values in (top) isolated cells, (middle) aggregates, and (bottom) monolayers. Units are scaled such that an MSD = 1 corresponds to the traversal of an area the size of an average simulated cell. (b) Plots of the diffusion coefficient across all J values for (top to bottom) isolated cells, aggregates, and monolayers. Diffusion coefficient units are scaled similarly to MSD in (a), where D = 1 corresponds to the cell-sized area that a typical cell will traverse in 1% of the total simulation time. (c) The power law diffusion exponents ⍺ (MSD ∝ t^α^) are plotted as a function of J value for (top to bottom) isolated cells, aggregates, and monolayers. The shaded regions in (b-c) for aggregates indicate where J_cell_ is sufficiently large relative to J_env_ such that fluctuations allow the eventual disintegration of the aggregate. Red dashed lines in (b-c) indicate the threshold for normal diffusion (⍺ ≥ 0.9) due to fluctuations of the anomalous diffusion coefficient around ⍺ = 1, as also described in Ref. 42. MSDs and related quantities used for fits are ensemble averaged over cells and simulations (n=20, single cell simulations; n=150, aggregate cells over 3 separate simulations; n=300, monolayers, over 3 separate simulations). Points highlighted by red asterisks denote behavior indicative of jamming for the interrogated metric. For diffusion coefficients (b), this reflects a D value sufficiently small such that cells traverse less than a cell-sized area over the entire simulation length, while for power law exponents (c) this indicates a sharp transition to sub-diffusion.

In aggregates, diffusion coefficients of cells gradually decrease with increasing J_cell_ until J_cell_ = J_env_, at which point there is an uptick due to the disintegration of the aggregate (**Fig. 2b, Video 1**). In contrast, trends in the anomalous diffusion exponent ⍺ are less straightforward, and the value of ⍺ depends on the magnitude of J_cell_ relative to J_env_. When J_cell_ = 0, cells are evidently very mobile (**Video 1**), as reflected in the extensive overlap of trajectories (**Fig. 1b**) and relatively large diffusion coefficients (**Fig. 2b**). However, the region cells can traverse is effectively bounded by the empty lattice due to the larger J_env_ parameter, resulting in saturation of the MSD. This leads to a power-law exponent consistent with sub-diffusion (**Fig. 2b**, ⍺ = 0.45). Such behavior is expected for diffusion within a bounded region^58^, although we do not observe saturation at a specific value, as global motion of the aggregate can and does occur at certain values of J_cell_ (**Video 1**, J_cell_ = 1). As J_cell_ is increased, cell mobility declines and the saturation of the MSD is not observed, leading to a larger value of the ⍺ parameter (**Fig. 2b**). Cells are super-diffusive (⍺ = 1.7) at J_cell_ = 1.75, where fluctuations lead to rapid detachment of a proportion of cells from the aggregate at long times (**Video 2**). When J_cell_ ≥ J_env_, cells are no longer confined by the environment and can freely diffuse throughout the lattice (⍺ ≈ 1).

Monolayers display distinct and monotonic changes of dynamics with varying J_cell_, owing to the simpler situation in which J_env_ plays no role. First, MSD curves span approximately 3 orders of magnitude over this range of J_cell_ values (**Fig. 2a**), with dramatic reductions in cell motility with increasing J_cell_ (**Video 1**). Unlike in aggregates, monolayers – lacking extracellular space – remain normally diffusive for J_cell_ values up to J_cell_ = 1 (**Fig. 2c**), after which evidence of anomalous sub-diffusion becomes obvious. Compared to aggregate simulations, monolayer simulations show a sharper drop in both anomalous diffusion exponent and diffusion coefficient (**Fig. 2b,c**). There is a striking decrease in D to near zero at J_cell_ = 1, while the transition from normal to sub-diffusion occurs slightly later, at J_cell_ = 1.25, transitioning from ⍺ ≈ 1 to ⍺ ≈ 0.4, deep in the sub-diffusive regime.

Given that varying the J parameters associated with cell adhesion alone can lead to a transition to sub-diffusive behavior in these simulations, even in the simple cases in which only J_env_ (isolated cells) or J_cell_ (monolayer) plays a role, we further examined the role of J values in effecting such behavior. The primary effect of increasing J is increasing energy associated with candidate spin flips and decreasing the fraction of spin flip acceptances during a given timestep (**Fig. S2**). As such, J acts as an effective temperature, though it is important to note that the CPM separately includes a temperature parameter that, when decreased, also results in decreased acceptance of spin flips. The temperature parameter is fixed at 1 in these simulations (**Table S1**), as temperature does not have a meaningful biological counterpart and thus variation of this parameter would not provide instructive results regarding cell dynamics. Directly comparing CPM simulations of isolated single cells and confluent monolayers, where J_env_ (relevant for isolated cells) = J_cell_ (relevant for cells in monolayer) across simulations, the effects of confinement become apparent. Despite identical adhesion penalties relative to monolayers, isolated cells remain normally diffusive over the full range of J values explored in this work (**Fig. 2c**), highlighting that the energy barrier imposed by a large J value alone is not sufficient to induce sub-diffusion. Closer inspection of diffusion coefficients in monolayers revealed indicators of glassy dynamics unique to this context. Diffusion coefficients of individual cells in monolayers were computed for individual cells by computing the time-averaged MSD for each cell trajectory. Single cell diffusion coefficients show log-normal distributions spanning several orders of magnitude (**Fig. S3**). Distributions of diffusion coefficients broaden with increasing J_cell_ in monolayers, a hallmark of glassy systems indicating heterogeneous dynamics^59^. Taken together, it is clear that crowding effects from monolayer confinement are critical to provoking sub-diffusive and frustrated, heterogeneous cell dynamics associated with glassy behavior and jamming.

### Cell Shapes in the Potts Model

Cell shapes are widely used as an indicator of jamming in a variety of experimental and simulated systems. Cell shape is typically defined as a dimensionless perimeter to area ratio (cell shape index = *P*/*A*^1/2^), which is a readily accessible quantity in both simulations and experimental work. The jamming transition is expected to occur at a shape index of 3.81, with values at or below 3.81 associated with jammed systems^34^. We considered cell shape index to assess whether this commonly used metric is associated with jamming in the CPM. To minimize grid effects in the CPM, cell perimeters were approximated using a previously described method (See **Supporting Text 1** for more detail)^46, 47^, and area was determined by counting the number of lattice sites comprising a given cell. For all simulations discussed so far, the target shape index was set to that of a circle (3.55), as shown in **Table S1**.

We first analyzed cell shapes by examining the fluctuations in shape over time (**Fig. S4**). In all cases, cell shape does not evolve appreciably after the early stages of the simulation; thus, we consider the distribution of cell shapes at the final timestep to be representative of the simulation as a whole. For aggregates, cell shape decreases with increasing J_cell_ but does not fully mirror the trends seen in **Fig. 2b**. Shape indices for simulations with J_cell_ = 0 – 0.75 lie above the expected jamming threshold, while shape indices for J_cell_ ≥ 1 fall below this cutoff index (**Fig. 3a**). In cases where aggregates disaggregate (J_cell_ ≥ 1.75, **Fig. 3a**), we do not expect cell shape to remain a useful structural metric of jamming, since the simulated cells are no longer in contact. Nevertheless, we show the complete set of shape indices for each simulated set. For aggregates, we also considered the possibility that the total number of cells might alter cell shape index, due to differences in cellular environment between interior and exterior cells, specifically in cases where J_cell_ < J_env_ and aggregates remain intact. However, we see no notable shift in cell shape indices as a function of aggregate size (**Fig. S5**), indicating negligible dependence on this variable. In monolayers, the shape index drops below the jamming threshold at J_cell_ = 1 (**Fig. 3b**), which closely corresponds with the observed transition to sub-diffusion, which occurs at J_cell_ = 1.25 (**Fig. 2c**). In addition, a narrowing of shape distributions at J_cell_ ≥ 1.25 is seen, consistent with the expectation that in deeply jammed states cell shapes become more uniform^37^.

**Figure 3.**
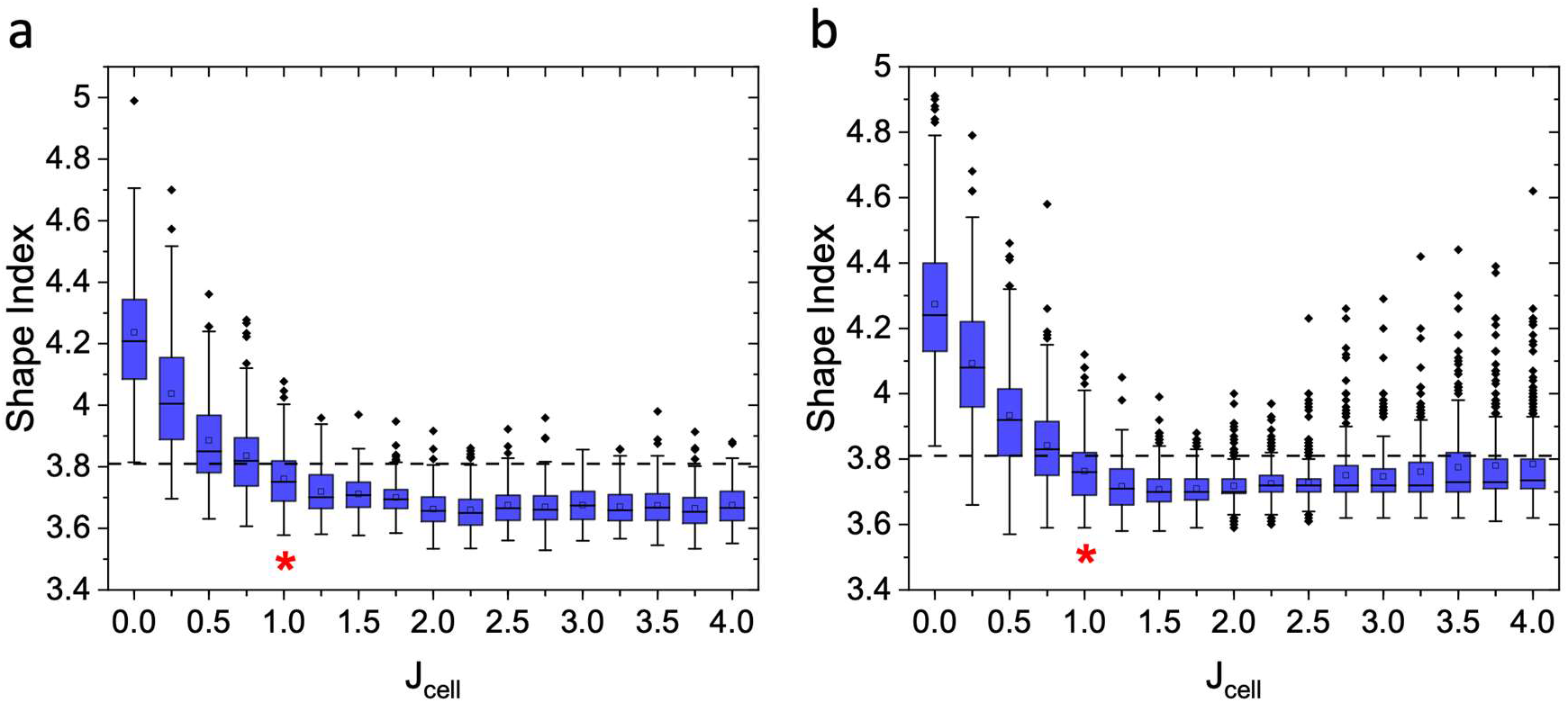
Comparison of cell shapes in CPM simulations. Final cell shapes in (a) aggregate and (b) monolayer simulations after 1×10^5^ MCS for each J_cell_ value. Shape index is defined as P/√A. The dashed horizontal line denotes the previously reported jamming threshold, at shape index of 3.81 as described in Ref. 34, with values below this threshold expected to jam. The blue boxed area denotes the interquartile range (IQR), whiskers denote 1.5*IQR, solid diamond points denote outliers, horizontal lines denote medians, and open squares denote means (aggregates: n=150, from 3 simulations; monolayers: n=300, from 3 simulations). Asterisks denote the values of J_cell_ at which mean/median shape indices dip below the expected jamming threshold.

### Cellular Rearrangements in the Potts Model

While reductions in cell motility, presence of sub-diffusive behavior, and changes in cell shape are all indicators of a jamming transition, we further analyzed cell behavior via quantification of cellular rearrangements. In densely packed tissue, cell motility will always be accompanied by neighbor exchange, though such exchange is limited in collective migration. In more commonly employed models of jamming, such as the vertex model, cellular rearrangements are explicitly included in the model. Specifically, in the absence of explicit inclusion of cell proliferation and cell death, cells can only rearrange via T1 transitions, a neighbor exchange involving four cells that consists of the disappearance of one cell junction in favor of the formation of another^60^. At the onset of jamming, as assessed through the shape index, T1 rearrangements cease, and cell motility is nearly abolished. In the CPM, the lattice-based nature and stochastic progression of the model makes the measurement of such rearrangements as well as their causal relationship with jamming less straightforward. We devised a metric for identifying neighbor exchanges as an analog to the T1 transition by measuring the changes in a cell’s local neighborhood (**Fig. 4**). Because of the site-by-site progression of spin flips in the CPM, cells experience frequent apparent neighbor changes via single or few lattice contacts with other cells, which are not representative of true cell rearrangements. Therefore, we considered the quantification of neighbor changes through two metrics. First, we counted the percentage of cells in the simulation that exchange any neighbors (even at the single lattice site level) at a variety of time intervals. Of cells experiencing neighbor changes, we next counted the fraction of these cells that only experience a change in a single neighboring cell during the given interval. Here, it is expected that as the time interval increases, the number of cells experiencing any neighbor exchange will increase, while the number of such cells experiencing a change in only one neighbor will decrease, as long as cells are exchanging neighbors beyond single lattice site contact fluctuations. Indeed, we find this behavior to hold for both aggregate and monolayer simulations (**Fig. 4**). We first interrogated the number of neighbor exchanges across time intervals spanning 100 – 3×10^4^ MCS in CPM aggregates (**Fig. 4a**). Here, we see the appearance of a crossover point, where the number of cells undergoing only single exchanges is exceeded by cells undergoing multiple changes of neighbors. Since the appearance of this crossover point occurs at the longest timescales, we next investigated the full range of J_cell_ at only the longest time interval (3×10^4^ MCS, **Fig. 4b**). The crossover point disappears at values of J_cell_ ≥ 1.25, which coincides with the J_cell_ values associated with minimum values for diffusion coefficients. Again, we note that due to aggregate dissolution when J_cell_ approaches J_env_, neighbor exchanges are no longer informative since cell-cell contacts are lost (**Fig. 4b**, shaded region). In monolayers, similar behavior is seen, with crossover points again observed at long measurement intervals for certain J_cell_ values (measurement interval 100 – 3×10^4^ MCS, **Fig. 4c**). As in aggregates, we examined the existence of crossover points over the full J_cell_ range (**Fig. 4d**). The disappearance of the crossover point occurs at J_cell_ = 1.25, coinciding with the transition from normal to sub-diffusion (**Fig. 2c**). For simulations with normally diffusive cells, the percentage of cells experiencing any neighbor changes approaches 100% at long measurement intervals, while the percentage of cells exchanging only a single neighbor approaches 0%. For simulations with cells showing sub-diffusive behavior, while the percentage of cells with neighbor exchanges does increase somewhat at long time intervals, the vast majority of these are single neighbor exchanges. Such single neighbor exchanges are not generally associated with cell motility, likely involving small changes in contacts caused by stochastic perimeter fluctuations. Importantly, the crossover metric described here accurately captures sub-diffusivity trends and does not misidentify contact fluctuations as functional cell motility.

**Figure 4.**
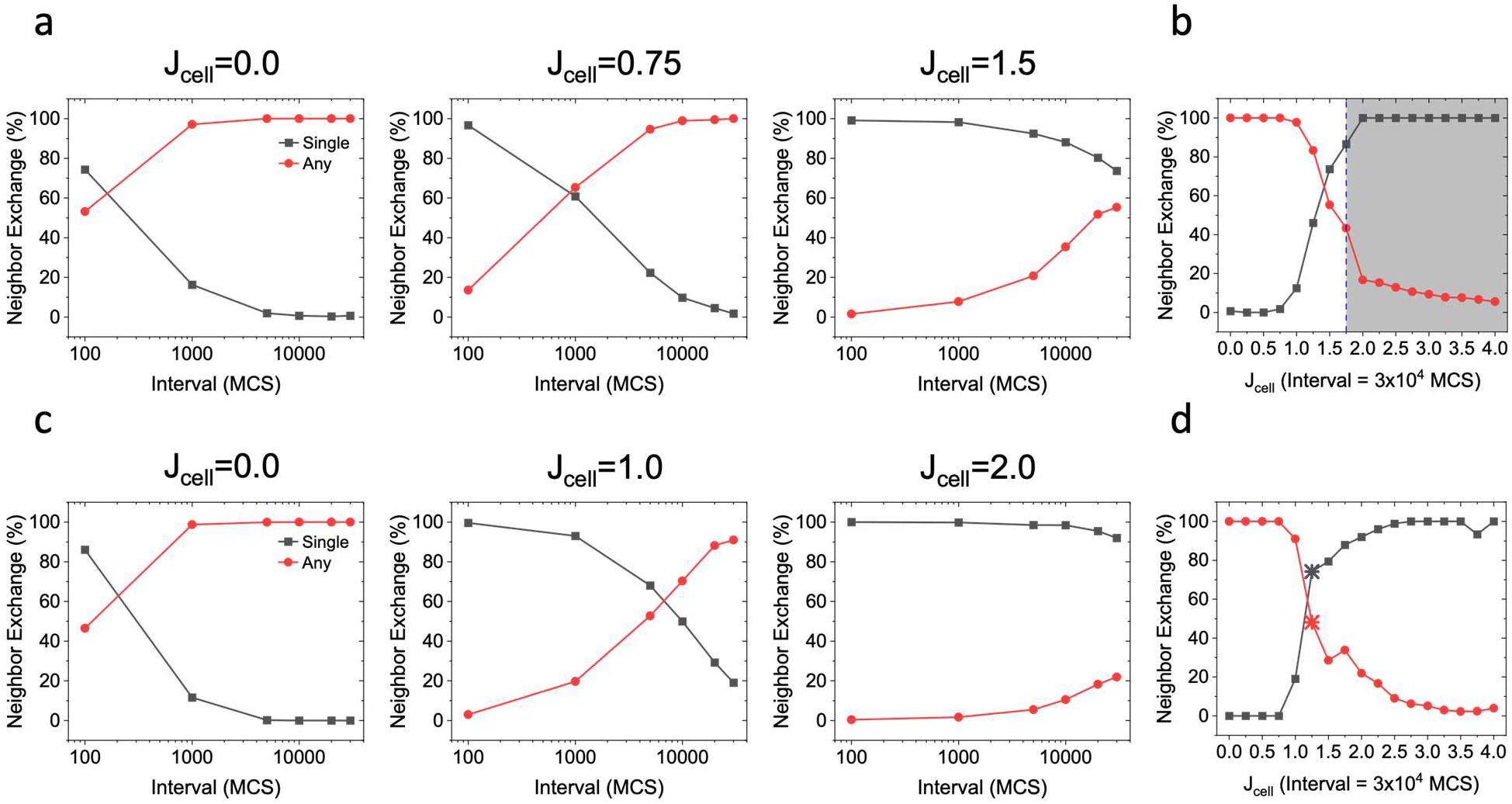
Neighbor exchanges in (a-b) aggregate and (c-d) monolayer simulations at different measurement intervals. “Any” neighbor exchange (red) represents the percentage of cells that experience any change in immediate neighborhood during the measurement interval. Among the cells that experience any neighbor exchange, “single” neighbor exchange (black curve) represents the fraction of this population that experiences a change in only one neighbor during the given measurement interval. (a,c) The measurement intervals span the range 100-3×10^4^ MCS for each J_cell_ value indicated, while (b,d) shows the percentages of each exchange type for all J_cell_ values at the maximum measurement interval in (a,c). The shaded region in (b) indicates the region where aggregate disintegration occurs. Asterisks indicate the values of J_cell_ following the crossover point, indicative of jamming.

### Effects of Cell Shape on Jamming in the CPM

We next varied target cell shape to probe the effects of shape on the onset of jamming within the CPM as well as to further explore the ability of the shape index to predict jamming in the CPM. As discussed above, the target area and target perimeter of cells were previously set to a circular target shape index of approximately 3.55. In experimental contexts, more diverse cell morphologies and, in particular, highly elongated cells are regularly observed, especially among aggressive cancer cells. Therefore, we altered target perimeter to probe the effect of shape variation on the fluid-like or solid-like behavior of cells in these simulations. Given the clearer signs of jamming in cellular monolayers relative to unconfined simulations, we restricted this analysis to monolayers where unjamming would be more challenging.

We hypothesized that increasing the target cell perimeter, while keeping all other variables constant, would increase cell shape index as well as cell membrane fluctuations, allowing cells to remain mobile and pushing the jamming transition to higher values of J_cell_, the adhesion parameter. Thus, we performed a set of monolayer simulations in which the target perimeter of cells was increased such that target shape index increased by a factor of ≈ 2 (**Table S1**), corresponding to more elongated cells. To evaluate the onset of jamming, again the diffusion coefficient, anomalous diffusion exponent, shape index, and neighbor exchanges were considered (**Fig. 5**). The first notable difference between these simulations and those with cells with a round target shape is a more gradual decrease in diffusion coefficient as J_cell_ increases (**Fig. 5a**), whereas previous monolayer simulations showed an abrupt drop in D associated with the onset of sub-diffusion (**Fig. 2b,c**). The onset of sub-diffusion here is not seen until J_cell_ = 2.75 (**Fig. 5b**). Like previous monolayer simulations (**Fig. 2c**), the drop from normal diffusion (⍺ ≈ 1) to sub-diffusion (⍺ ≈ 0.4) is abrupt. A steady decrease in observed cell shape index is seen as a function of increasing J_cell_ (**Fig. 5c**), approaching the jamming threshold of 3.81. As in cell shape quantification in earlier simulations (**Fig. 3**), the onset of sub-diffusion does not perfectly correspond to the expected jamming threshold as assessed by cell shape index – however, there is a slight drop and then plateau in cell shape index at J_cell_ ≥ 2.75. The appearance of the crossover point in neighbor exchanges shifts to nearly match the J_cell_ value associated with the onset of sub-diffusion, with significant neighbor exchanges resulting in crossover at all J_cell_ < 2.5 (**Fig. 5d**). Taken together, these metrics demonstrate that the increase in target cell shape index pushes the onset of jamming to high values of J_cell_ associated with significant penalties for disrupting initial cell-cell contacts. The shift of this transition can be further rationalized by considering the effect of target perimeter variation on the interaction between cells. As in other work^46^, we can define an interfacial tension, γ, by taking the derivative of the CPM Hamiltonian with respect to perimeter:

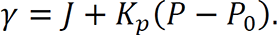

(For a detailed discussion of this equation, see **Supporting Text 2**). For round cells (target shape index 3.55), the second term of this equation is approximately 0, as target perimeter is effectively reached in all cases, and the interfacial tension is given simply by γ ≈ J. However, for cells with a larger target perimeter, the second term is typically negative, lowering the interfacial tension between cells to values that permit cells to remain motile. Examining the average interfacial tension per cell across both elongated and round cells reveals that there is a common value for interfacial tension associated with the onset of sub-diffusion (**Fig. 5e**).

**Figure 5.**
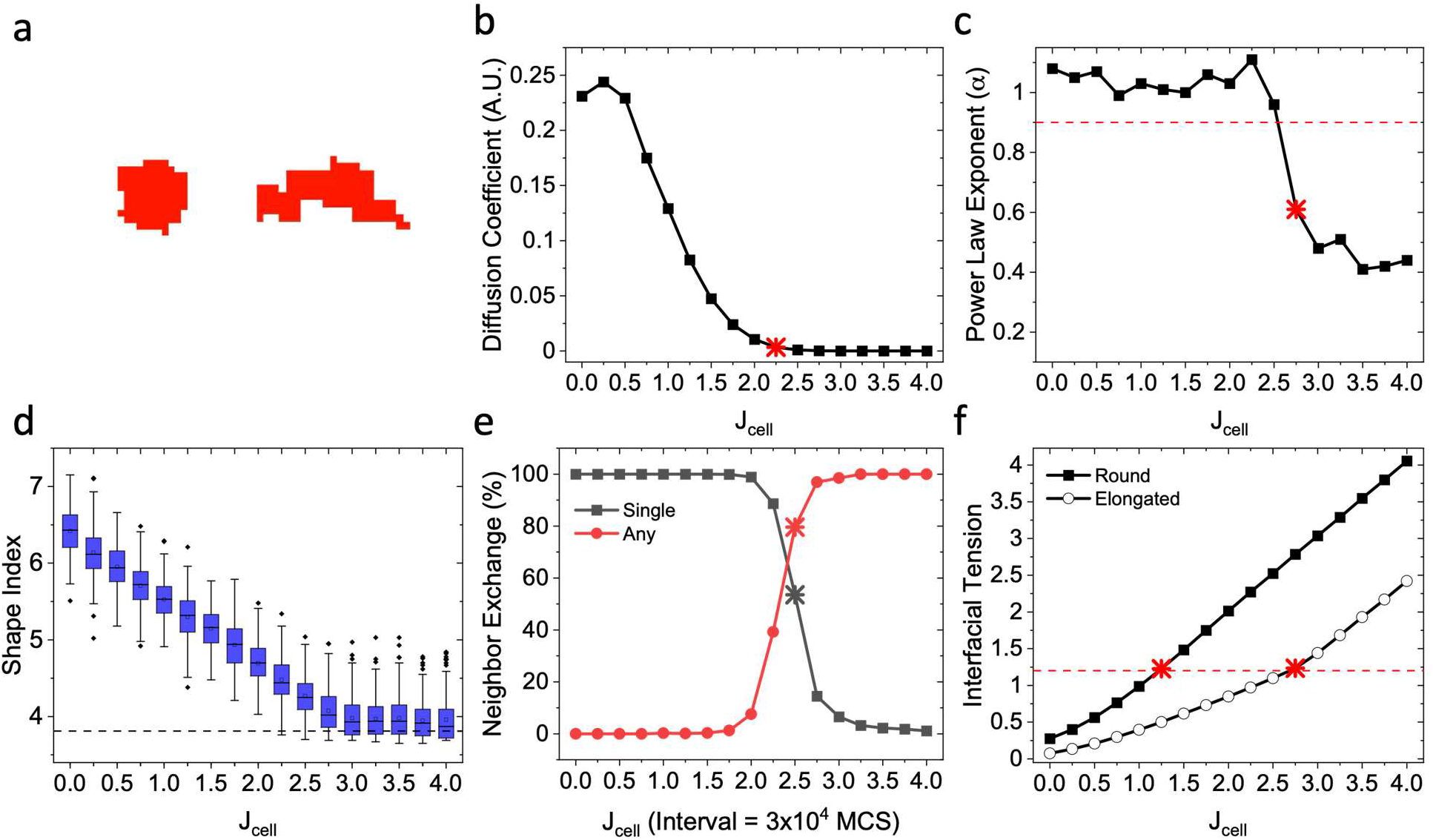
Jamming metrics for a monolayer of elongated cells. (a) Representative (left) elongated cell and (right) round cell, with elongated cells having a shape index approximately 2 times greater than round cells. Both cell types are depicted at J_cell_ = 1 after equilibration of the CPM monolayer, with target perimeters of 70.5 and 35.5, respectively. (b) Diffusion coefficients for all J_cell_ values displayed in the same units as Fig. 2. (c) Anomalous diffusion exponents from power law fits as a function of J_cell_ for the monolayer. The dashed line indicates the threshold for normal diffusion (⍺ = 0.9) as explained in Fig. 2. (d) Cell shape indices for the final state as a function of J_cell_, with the dashed line indicating the expected jamming threshold as described in Fig. 3. (e) Neighbor exchange analysis of monolayers using the same metrics described in Fig. 4, interrogated for all J_cell_ values at a measurement interval of 3×10^4^ MCS. (f) Interfacial tension in round and elongated monolayer simulations, as defined in the text. Dotted red line denotes the value associated with a transition to sub-diffusion, which occurs at J_cell_ = 1.25 and 2.50 for round and elongated cells in monolayer, respectively. As in prior figures, asterisks denote behavior consistent with jamming as indicated by each metric.

To further characterize the dynamics of CPM monolayers, we more closely examined the cell migration seen in such monolayers. In round cells, it is apparent that the transition from mobile to jammed cells occurs over a relatively narrow range of J_cell_ (**Video 3**). Moreover, at low J_cell_ values, cells display individual cell migration, while migration becomes increasingly collective as the jamming transition is approached. To confirm this observation, we calculated spatial correlations in cell velocities at a variety of J_cell_ values. Spatial correlations in velocity were determined according to,

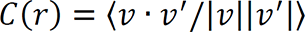

where v and v’ denote velocity vectors of different cells, r is the distance between cells, and brackets denote averaging over all lag times of 1000 MCS and all pairs of cells. In monolayers consisting of round cells, an increase in spatial correlation between neighboring cells is observed as the jamming transition is approached (**Fig. 6a**). Spatial correlations vary in accordance with the average distance between neighboring cells and become longer range in the vicinity of the jamming transition, indicating collective cell motion. The correlation is maximized at J_cell_ = 1, closely matching the adhesion parameter value associated with the sharp drop in diffusion coefficient and abrupt transition to sub-diffusion (**Fig. 2b,c**). At higher J_cell_ values, spatial correlations in velocity decrease as the system moves deeper into the jammed state (**Fig. 6a**). For cells with a larger target cell shape index, no spatial correlations in velocity are seen in the vicinity of the jamming transition nor are any strong correlations found across the range of J_cell_ values explored (**Fig. 6b**). Comparison of round and elongated cell monolayers at J_cell_ = 2 (**Video 4**) reveals clear differences in migratory character, where elongated cells display a considerable motility increase and absence of collective migration relative to round cells of the same J_cell_. Furthermore, elongated cells do not show collective motility over the broader J_cell_ range (**Video 5**), suggesting that such cells do not undergo the same transitions from single cell motility to collective migration to the jammed state, as seen in round cells, and that collective migration does not necessarily precede jamming.

**Figure 6.**
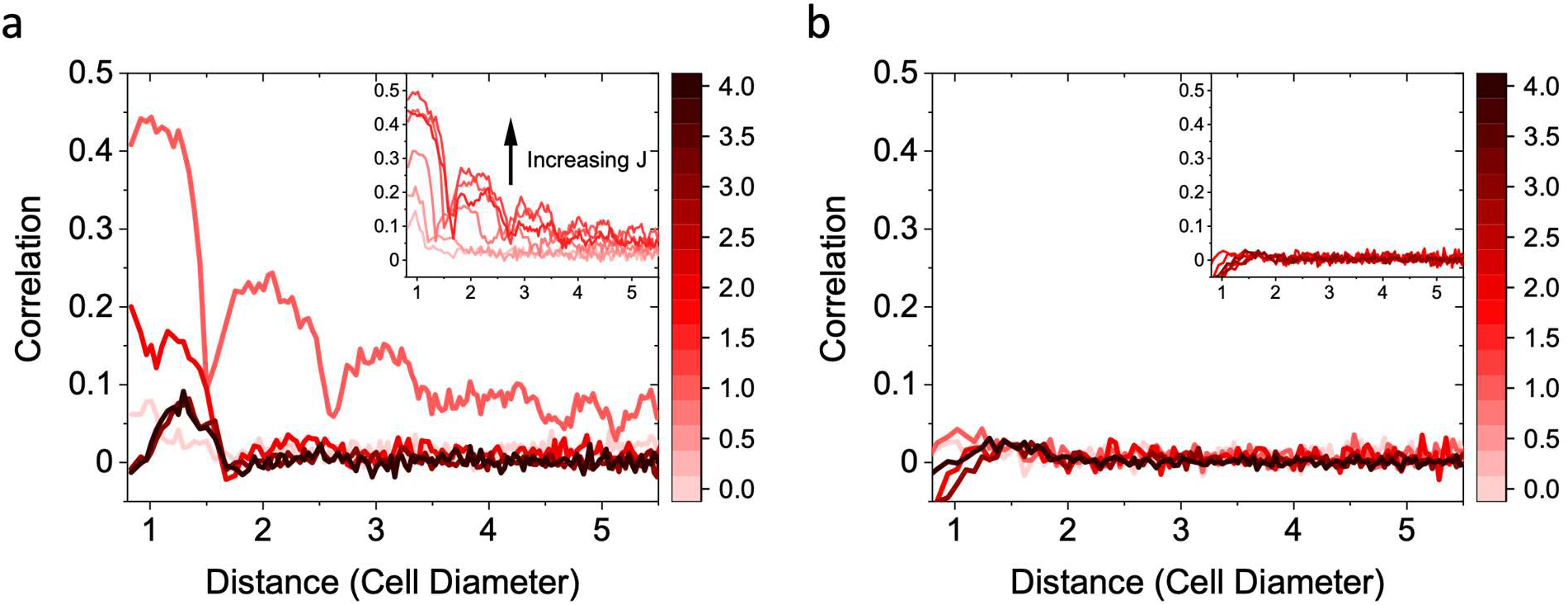
Spatial correlations in velocity indicative of collective motion in monolayers. (a) Spatial correlations in round cell monolayers for J_cell_ = 0,1,2,3,4 and (inset) near the onset of sub-diffusion (J_cell_ = 0.25, 0.50, 0.75, 1.00, 1.25, 1.50). (b) Spatial correlations in elongated cell monolayers over the complete J_cell_ range (J_cell_ = 0, 1, 2, 3, 4) and (inset) a smaller range near the onset of sub-diffusion (J_cell_ = 1.50, 1.75, 2.00, 2.25, 2.75, 3.00). Curves were smoothed with a moving average. Colorbar shows J values.

### Effects of Shape Heterogeneity on Jamming in the CPM

To investigate one of the possible effects of EMT induction within a jammed monolayer, we examined the effect of mixing cells with target shape indices associated with elongated and round cells in a single monolayer to probe whether cell elongation – decoupled from an explicit change in adhesion energy – was capable of fluidizing otherwise jammed rounder cells. We combined round cells with adhesion parameter just past the jamming threshold (J_cell_ =1.50, which yielded ⍺ = 0.4 for the anomalous diffusion exponent for single component monolayers of this type), representative of jammed epithelial cells, with elongated cells (also with J_cell_ = 1.50), representative of mesenchymal cells, in a 1:1 ratio and simulated the combination for 1 × 10^5^ MCS. Indeed, the mixed monolayer remained fluid, as indicated by larger diffusion coefficients relative to round cell monolayers (**Fig. 7a**, **Fig. 2b**) an anomalous diffusion exponent near 1 (**Fig. 7b**, ⍺_elongated_ = 1.01, ⍺_round_ = 0.96, ⍺_all_ = 1.01), indicative of normal diffusion. Decreasing the proportion of elongated cells still resulted in relatively effective fluidization of the monolayer. Monolayers with as few as 33% of cells having the higher target cell shape index are normally diffusive (⍺_all_ = 0.92), and those with 25% elongated cells remains nearly normally diffusive as well (⍺_all_ = 0.87). Even at the smallest fraction of elongated cells explored here (20%), cells are only mildly sub-diffusive, distinct from the deeply sub-diffusive behavior observed in cell monolayers in which the target shape is a circle.

**Figure 7.**
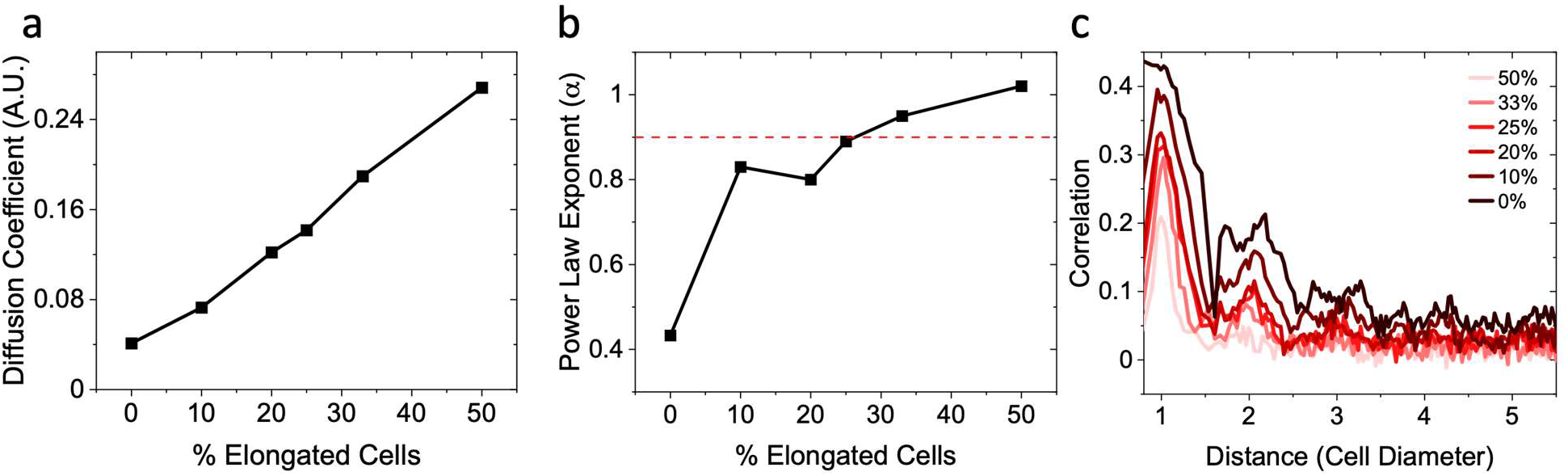
Diffusive character and spatial correlations in mixed monolayers composed of round (target shape = 3.55) and elongated (target shape = 7.05) cells with the same J_cell_ = 1.50. (a) Diffusion coefficients and (b) anomalous diffusion parameter, ⍺, for mixed monolayers of increasing elongated cell composition. Diffusion coefficients are expressed in the same scaled units as described in Fig. 2. (c) Spatial velocity correlations in monolayers as a function of elongated cell composition, for the same J_cell_ = 1.50 in (a,b).

To further probe cell dynamics in these mixed monolayers, spatial correlations in cell velocities were assessed. Correlations in velocities decrease across the monolayer with increasing percentage of elongated cells (**Fig. 7c**). In a mixed monolayer, strong correlations are only observed for nearest neighbor cells, with weak correlations observed for next nearest neighbors at lower percentages of elongated cells. At all elongated cell percentages, correlations in space more quickly decay to zero relative to the purely round cell case. It is apparent that the ability of cells to elongate inhibits jamming while decreasing the degree of correlated motion, abolishing the apparent transition to collective motion that precedes the jamming transition.

## Discussion

The work presented here demonstrates that cellular Potts model simulations show jamming that depends on cellular adhesion energies, degree of cellular confinement, and cell shape. The model systems explored in this work - single cells, aggregates and monolayers - represent three fundamentally different types of environments. Cells in isolation experience energy barriers dictated by adhesion energy associated with cell-environment contacts (J_env_), but no confinement or physical barriers to movement. Cells in aggregates experience a degree of confinement from surrounding cells and, when J_cell_ << J_env_, the surrounding environmental space. Although the environment can contribute to cell confinement, empty space around the aggregate can also provide additional freedom for cell motility and lead to lower overall cell density in the aggregate relative to a confluent monolayer. In a monolayer, confinement is much more severe, and cells must navigate the densely packed environment to remain mobile. We find that confinement of cells in such monolayers leads to abrupt changes in cell motility as a function of adhesion energy, leading to behavior reminiscent of a jamming transition. Importantly, the metrics used here are in good agreement with each other in regard to the value of the cell adhesion parameter at which jamming occurs in monolayers (**Table S2**), supporting the existence of a jamming transition in this context.

The effects of confinement are reflected in the magnitude and character of mean square displacements in each system (**Fig. 2**). Isolated cells remain normally diffusive at all adhesion energies (**Fig. 2a**), while aggregate cell sub-diffusion is dependent on the strength of environmental confinement (**Fig. 2b**). In contrast, monolayers experience an abrupt transition to deeply sub-diffusive behavior and multiple orders of magnitude decrease in diffusion coefficients across this same range of J_cell_ values (**Fig. 2c**). These differences highlight that abrupt cell arrest is unique to monolayers and emphasizes how degree of confinement affects cell dynamics. Importantly, sub-diffusion and arrest are not solely linked to energy barriers dictated by the adhesion parameter J, as shown by the normally diffusive character of isolated cell migration.

Sub-diffusion has previously been used as the primary readout for the onset of jamming within the CPM^42^, and here we again find it to be a generally reliable indicator in the confluent monolayer. However, in the set of simulations described here, we employed additional metrics to characterize the onset of the jamming transition to provide a fuller framework for analysis of CPM-based jamming, as well as avoid mischaracterization of situations in which sub-diffusion is not associated with jamming, as observed in aggregates. We considered not only the onset of sub-diffusion but also cell shapes, cellular rearrangements, and spatial velocity correlations. These metrics support the existence of jamming-induced sub-diffusion in monolayers and were generally in excellent agreement as predictors of the jamming transition (**Table S2**). Equilibrium cell shape in the CPM is moderately dependent on the value of J_cell_, as cells tend to alter their perimeter to either minimize or maximize contacts with the surroundings. Therefore, as adhesion penalty is increased, a decrease in cell shape index occurs as cells reduce contact length, pushing the cells towards jamming. Shape index distributions follow trends seen in other systems, with the shape index of cells with a round target shape reaching and dipping below the previously identified jamming-associated shape index of 3.81 at the onset of sub-diffusive behavior (**Fig. 3a**)^34^. Simulations of cells with a higher target shape index, elongated cells, do not show perfect correspondence between the onset of jamming and cell shape index dipping below 3.81 (**Fig. 3b**), though trends in average shape index and distribution are consistent with previous findings, with a narrowing distribution and stabilization of the mean cell shape index at a value of ∼3.9 for cells that exhibit other features associated with jamming.

While cell shape analysis did not perfectly conform to expected thresholds of jamming identified in other systems, trends in cell shape were in line with previous results. As expected, increasing cell shape index via an increase in the target perimeter resulted in very effective fluidization of otherwise jammed cells (**Fig. 5**). Although jamming in the CPM is highly dependent on cell-cell adhesion as set by J_cell_, as shown in this work and others^42^, increase in target perimeter (without any change in J values) provides cells with sufficient degrees of freedom to overcome otherwise prohibitively high energy barriers associated with cell-cell rearrangements in the monolayer. Through neighbor exchange analysis, we provided a clear measure of cell rearrangements within the CPM and confirmed that the absence of such rearrangements accompanies the abrupt transition to sub-diffusion that was observed (**Fig. 5e**). Notably, there is a common interfacial tension associated with jamming that applies across cells of different morphologies (**Fig. 5f**). Such a universal descriptor of jamming could be used to predict jamming across a variety of contexts not limited to the CPM and opens the door for further comparison to experimental data and other *in silico* models of epithelial systems.

Collective migration is inherently intertwined with the jamming transition^10, 18, 27^, in that it allows cells to continue to remain mobile when local caging sets in and energy costs to neighbor exchanges become high^61–63^. In this work, such motility manifests in migration of round cells in epithelial monolayers (**Video 3, Fig. 6a**). Spatial velocity correlations provide information on collective cell dynamics within the CPM systems studied here. Velocity correlations in monolayer simulations of round cells reveal three distinct regimes of cell behavior as a function of the cell adhesion parameter. At very low J_cell_ values, cell motion is disordered and minimal correlations in velocity are present – presumably, cells are free to explore the space with minimal barriers to rearrangement. As described in prior vertex model studies, energy barriers to cell rearrangements effectively vanish in this regime^34^. As the jamming transition is approached (upon increasing J_cell_), collective cell migration emerges, and long-range correlations in velocities that extend over multiple cell diameters are observed. It is particularly interesting that this signature of collective cell migration occurs in the minimal implementation of the CPM utilized in this work. In other iterations of the CPM where collective migration has been observed, the model includes an explicit cell motility term in the Hamiltonian that encourages directed cell motion and results in collective migration^50, 51^. In contrast, in this work, there is no explicit cell motility term, and collective cell motion arises naturally near the onset of the jamming transition. In this regime, cell rearrangements become rare, and cell motion can only be achieved through collective flow of the monolayer. The third and final regime occurs upon complete arrest of cells, at high J_cell_ values, such that cell motility that requires new cell contacts becomes sufficiently unfavorable. Here the absence of velocity correlations in combination with the cessation of cell rearrangements demonstrates that neither cell neighbor exchange nor collective flowing of the monolayer is permitted. Cell shape fluctuations among cells with constant J_cell_ adhesion penalties allow cells to avoid collective migration entirely, demonstrating that collective migration prior to jamming is not observed for all cell types.

The results presented in this work highlight the ways in which cell shape influences the onset of jamming. When forcing cell elongation through an increase in cell target perimeter, a significant shift in the onset of jamming as a function of adhesion energy was observed. It is important to note that, while the onset of jamming is clearly pushed to higher values of J_cell_, encoding a large target perimeter does not lead to a large increase in magnitudes of monolayer MSDs or diffusion coefficients. As can be seen in **Figs. 2-4**, the diffusion coefficients are very similar across elongated and round cell type monolayers of a given J_cell_ value. Rather, the main difference between the two cases is seen in the velocity correlations. Velocity correlations are effectively abolished in the monolayer with cells of higher target cell shape index, which gives cells the ability to deform (**Fig. 6b**). This same effect is seen in the mixed monolayers, where even relatively small percentages of elongated cells in the monolayer significantly reduces long-range velocity correlations (**Fig. 7c**). Unlike rounder cells, monolayers containing elongated cells display greatly reduced collective dynamics, maintaining more disordered motion until the onset of complete arrest at very high J_cell_ values, as they retain sufficient freedom of mobility to not have to rely on collective motion.

The results presented here have implications for the study of metastasis, particularly when considering cancer invasion as an unjamming transition. Cancerous tumors contain a diverse mixture of cell types, and the complex interplay between cells in this heterogeneous environment can influence the occurrence of jamming and unjamming transitions. Prior works have considered the interactions between potentially invasive, mesenchymal cells, and non-invasive epithelial cells, which will have tendencies towards unjamming and jamming, respectively^16^. Notable features of invasive cancer cells, which have progressed through the EMT, are elongated morphologies and softer, more deformable cell bodies. It is well-documented that transformed cells are consistently softer than healthy cells, which aids in their invasion by facilitating more efficient cell shape change that allows squeezing around obstacles and/or through tight spaces^47, 64–66^. Our results on elongated cells within the CPM suggest one way in which a deformable cell body can facilitate motility and invasion in highly crowded environments such as tissue.

## Acknowledgements

L.K. was supported by the Amgen Scholars Program and D.J.L. was supported by the Rabi Scholars Program. We thank Esha Maharishi for her contributions to CPM-based projects in the lab that led to this work.

## Data Availability

Python/C++ code used to initialize and run the CPM as well as to visualize the output is provided at https://github.com/kaufmanlab-columbia/CellularPottsModel.

## Author Contributions

Conceptualization: A.J.D., L.J.K.; Methodology: A.J.D., L.J.K.; Model Implementation: A.J.D., D.J.L., L.K.; Formal analysis: A.J.D., L.J.K.; Writing - original draft: A.J.D.; Writing - review & editing: A.J.D., L.J.K.; Data Visualization: A.J.D., L.K., L.J.K.; Supervision: L.J.K.; Project administration: L.J.K.; Funding acquisition: L.J.K.

