## Supporting Information for "Signatures of Jamming in the Cellular Potts Model"

### Supporting Text 1

### Detailed Analysis of Cell Shapes in the CPM

As noted in other work, perimeters in the CPM suffer from systematic and significant overcounting that originates from the underlying square lattice. Specifically, when measuring the perimeter by counting all lattice site edges, which we denote as the perimeter type $P_{CPM,}$ grid effects artificially inflate perimeters and therefore cell shape indices[1–3]. Given the importance of cell shape as a structural predictor of jamming, the ability to translate cell shapes to systems outside of the Potts model is especially critical for analysis of jamming transitions. For this reason, a perimeter approximation is preferred, such that cell perimeters in the CPM can be mapped to experimentally meaningful cell perimeters and/or cells in other models of cell migration (for example, vertex models).

After running CPM simulations, for the most accurate determination of equilibrium cell shapes (one that does not overestimate geometric perimeter because of the lattice nature of simulation), we utilized the method described in Ref. 1. Here, cell perimeter is determined by counting all non-cell sites within a specified neighborhood radius for each lattice site belonging to a cell, and then dividing this total perimeter count by an empirically determined correction factor to yield the geometric perimeter. In general, larger search radii provide a more accurate estimation of the geometric perimeter, although the search radius used should be appropriately matched to the size of the CPM cells. After exploring several different measurement configurations from Ref. 1, we found that a search radius of √13 (and associated correction factor of 36) was optimal for approximating perimeter in the system used in this study. For clarity, we term perimeters measured via this method $P_{true}$, in contrast to $P_{CPM}$.

While the correction method described above is effective in avoiding lattice artifacts and reporting accurate cell shapes, it is computationally expensive to implement during simulations. Therefore, we only utilize it to compute equilibrium cell shapes from the final configurations, such that the shapes obtained are comparable outside of the CPM framework. Within the CPM simulation, perimeter is calculated on the fly using a simplified assessment, approximating perimeter by counting lattice sites (rather than lattice site edges) located at the cell boundary as 1 perimeter site, and perimeters determined via this method are denoted $P_{app}$. This method avoids the lattice counting artifact but does introduce systematic error into the measurement of cell shapes. Nevertheless, $P_{app}$ reports similar behavior to $P_{true}$ as function of $J_{cell}$, notably much more reliably than $P_{CPM}$.

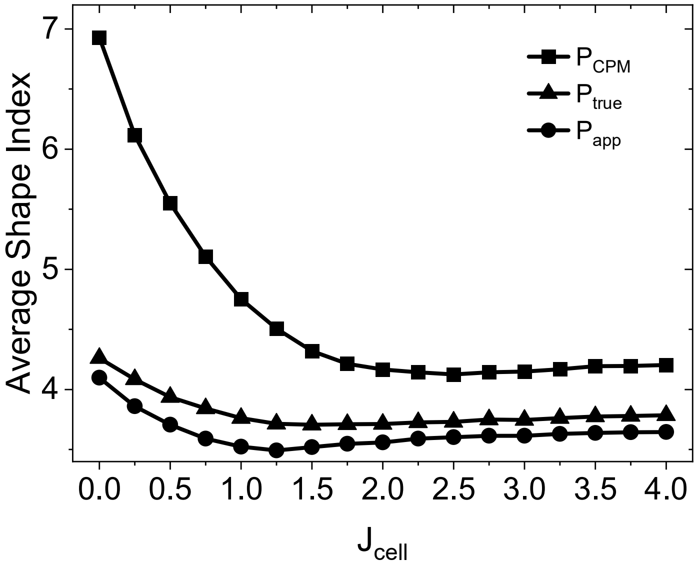


**Fig. ST1:** Mean cell shape indices as a function of $J_{cell}$ for round cell monolayers, determined via the different perimeter measurement methods described in **Supporting Text 1**.

**Supporting Text 2**

**Interfacial Tension in the CPM**

Interfacial tension ($\gamma$) is defined as the derivative of the total energy with respect to perimeter as described in Ref. 1. Given that only the adhesion and perimeter components of the energy depend on cell perimeter, it follows that:

$$\mathrm{H} = \mathrm{H}_{\mathrm{adhesion}} + \mathrm{H}_{\mathrm{perimeter}} + \mathrm{H}_{\mathrm{area}}$$

$$\gamma = \frac{\partial \mathrm{H}}{\partial \mathrm{P}} = \frac{\partial \mathrm{H}_{\mathrm{adhesion}}}{\partial \mathrm{P}} + \frac{\partial \mathrm{H}_{\mathrm{perimeter}}}{\partial \mathrm{P}}$$

In our implementation of the CPM, the measurement of perimeter differs in the adhesion and perimeter components of the total energy. Typically, the adhesion term is calculated by summing neighbor contributions over all lattice sites either within the von Neumann or Moore neighborhood (here we use von Neumann neighborhood). For a single J interaction energy, as is the case in monolayers, this yields:

$$\mathrm{H}_{\mathrm{adhesion}} = \sum_{(\mathrm{i,j})(\mathrm{i}',\mathrm{j}')\mathrm{neighbors}} \mathrm{J}(1 - \delta_{\sigma(\mathrm{i,j}),\sigma(\mathrm{i}',\mathrm{j}')})$$

Using the perimeter definitions from **Supporting Text 1**, this expression can be simplified to:

$$\mathrm{H}_{\mathrm{adhesion}} = \textstyle\sum_{\sigma} \mathrm{J} \cdot \mathrm{P}_{\mathrm{CPM}(\sigma)},$$

where we express the perimeter energy as a sum over cells using $\mathrm{P}_{\mathrm{CPM}}$, which reflects the perimeter from assessing all lattice neighbors within the von Neumann neighborhood at the cell boundary sites. Recalling that in the CPM Hamiltonian the perimeter energy term uses a different perimeter metric, $\mathrm{P}_{\mathrm{app}}$, this results in two different perimeter measurements appearing in the Hamiltonian during CPM simulations, via

$$\mathrm{H} = \sum_{\sigma} \mathrm{J} \cdot \mathrm{P}_{\mathrm{CPM}(\sigma)} + \mathrm{K}_{\mathrm{P}} \left(\mathrm{P}_{\mathrm{app}(\sigma)} - \mathrm{P}_0\right)^2 + \mathrm{H}_{\mathrm{area}}$$

To illustrate the differences in these perimeter measurements, we consider the following portion of a cell in the CPM lattice:

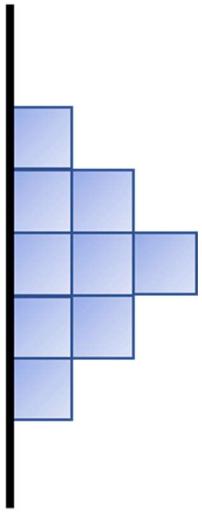

Here, $P_{CPM} = 11$, while $P_{app} = 5$. Rewriting the expression for interfacial tension with $P_{CPM}$ and $P_{app}$ gives:

$$\gamma = \frac{\partial H_{adhesion}}{\partial P_{CPM}} + \frac{\partial H_{perimeter}}{\partial P_{app}} \cdot \frac{\partial P_{app}}{\partial P_{CPM}}$$

Consider the spin flip depicted below, in which the cell gains a lattice site at the leading edge. Given the stepwise and discrete nature of the CPM, a single spin flip is the unit of perimeter growth that a cell undergoes at any given time point:

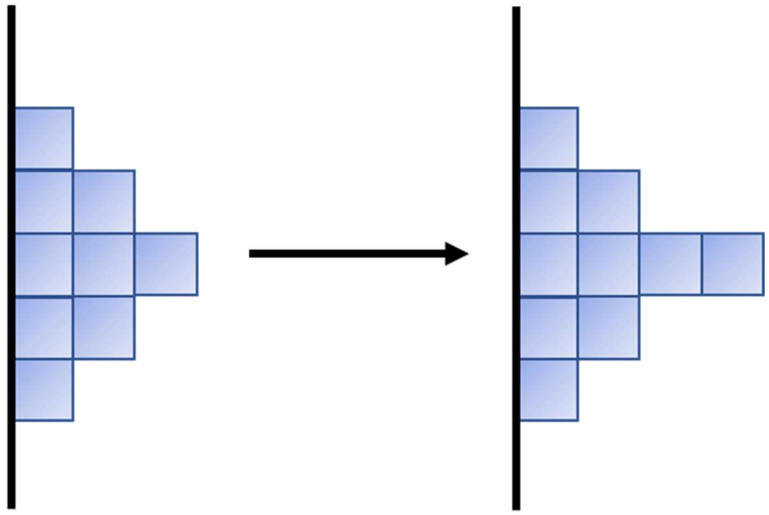

This cell has $P_{CPM(initial)} = 11$, and $P_{CPM(final)} = 13$, while by the approximated metric $P_{app(initial)} = 5$ and $P_{app(final)} = 6$, yielding $\Delta P_{CPM} = 2$ and $\Delta P_{app} = 1$. The interfacial tension of the cell-cell interface can then be approximated as follows:

$$\gamma \approx \frac{\partial H_{adhesion}}{\partial P_{CPM}} + \frac{\partial H_{perimeter}}{\partial P_{app}} \cdot \frac{\Delta P_{app}}{\Delta P_{CPM}}$$

$$\gamma \approx J_{cell} + 2K_P\left(P_{app} - P_0\right) \cdot \frac{1}{2}$$

$$\gamma \approx J_{cell} + K_P\left(P_{app} - P_0\right)$$

While the exemplary spin flip shows one of several possible spin flip scenarios, this relationship will also generally hold for other possible spin flips in this implementation of the CPM.

### Supporting Tables and Figures

#### Table S1

| | Isolated | Aggregate | Monolayer |
|---|---|---|---|
| Temperature (T) | 1 | 1 | 1 |
| Lattice Length (L) | 100 | 100 | 100 |
| Target Area ($A_{\tau(\sigma)}$) | 100 | 100 | 100 |
| Target Perimeter (Round, $P_{\tau(\sigma)}$) | 35.5 [3.55] | 35.5 [3.55] | 35.5 [3.55] |
| Target Perimeter (Elongated, $P_{\tau(\sigma)}$) | N/A | N/A | 70.5 [7.05] |
| Area Constant ($K_A$) | 0.60 | 0.60 | 0.60 |
| Perimeter Constant ($K_P$) | 0.05 | 0.05 | 0.05 |
| $J_{env}$ | variable | 2 | N/A |
| $J_{cell}$ | N/A | variable | variable |

**Table S1.** CPM parameters for each simulation type. T, $K_A$, and $K_P$ are held constant across all simulations. N/A indicates simulation parameters that were not used for a given case, while "variable" indicates parameters that are varied, as described in the main text. Square brackets ([]) denote the target shape index for cells with the perimeter and area indicated within.

**Table S2**

| | Round | Elongated |
|---|---|---|
| Diffusion Coefficient | 1.25 | 2.25 |
| Sub-Diffusion | 1.25 | 2.75 |
| Shape Index | 1.00 | > 4.00 |
| Neighbor Exchange | 1.25 | 2.50 |
| Interfacial Tension | 1.25* | 2.75* |

**Table S2.** Quantification of cellular jamming in monolayers via the different metrics explored in this work. The $J_{cell}$ values associated with jamming for round cell and elongated cell monolayers are depicted in the table. (*) For interfacial tensions, while a critical tension has not been explicitly associated with the jamming transition previously, coincident interfacial tension occurs at the $J_{cell}$ values listed.

**Figure S1**

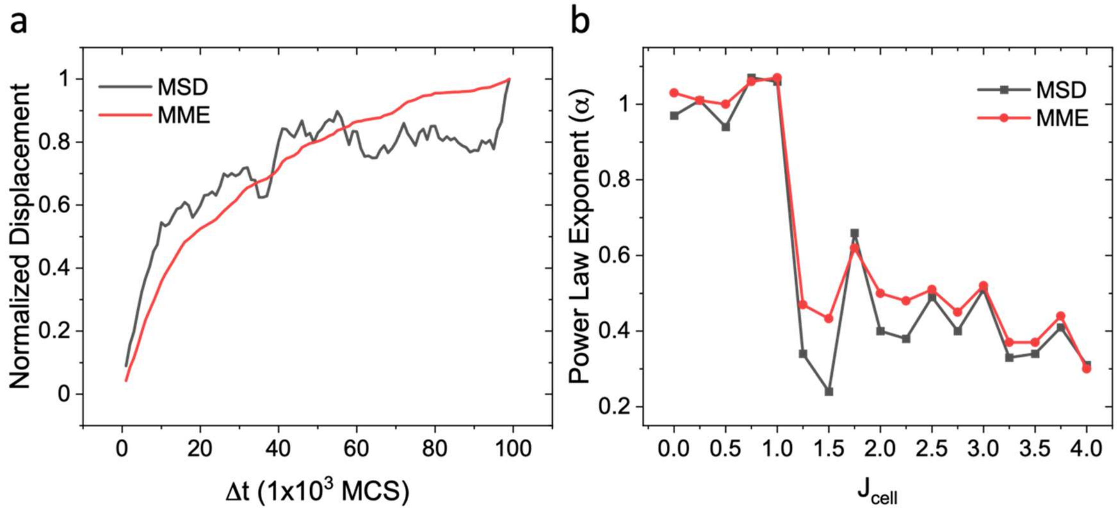


**Figure S1.** Comparison of displacement curves computed from ensemble averaged mean squared displacement (MSD) and ensemble averaged mean-maximal excursion (MME). (a) MME curves are considerably less noisy than MSD curves and give better quality fits (shown for aggregate, $J_{cell}$ = 0). (b) MME and MSD give similar power-law exponents describing the character of the diffusion (shown for round cell monolayers).

**Figure S2**

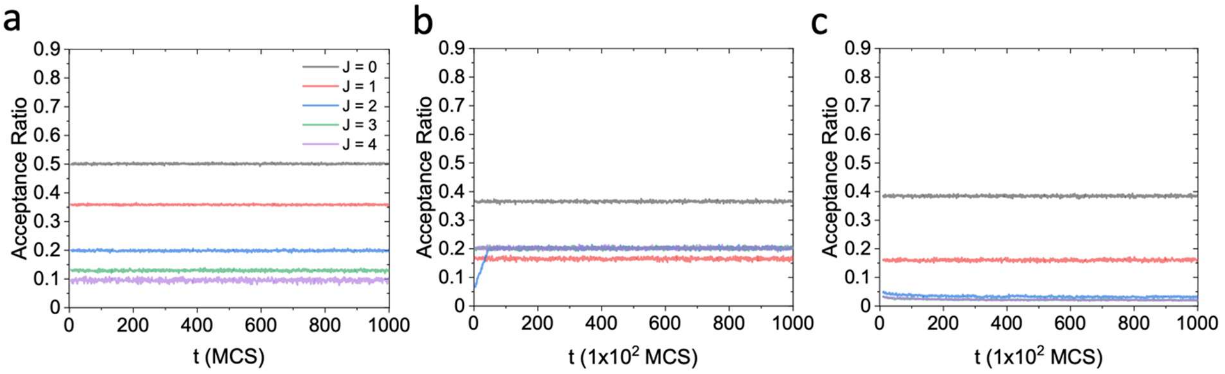


**Figure S2.** Acceptance ratios in (a) single cell, (b) aggregate, and (c) monolayer CPM simulations. The legend in (a) also applies to (b-c), where J denotes the relevant J parameter for each simulation type, as described in the main text.

**Figure S3**

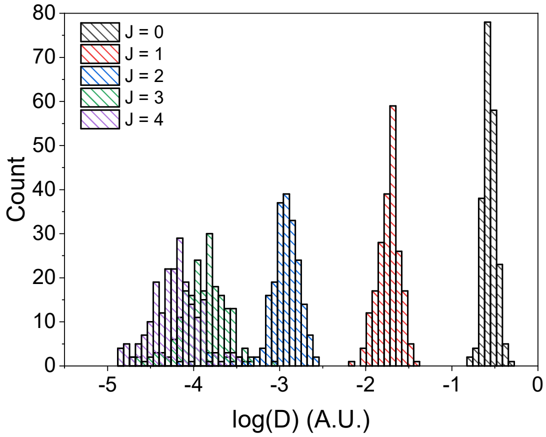


**Figure S3.** Diffusion coefficients of individual cells computed from time-averaged MSDs of single trajectories. Distributions of diffusion coefficients are shown for monolayer simulations as a function of $J_{cell}$.

**Figure S4**

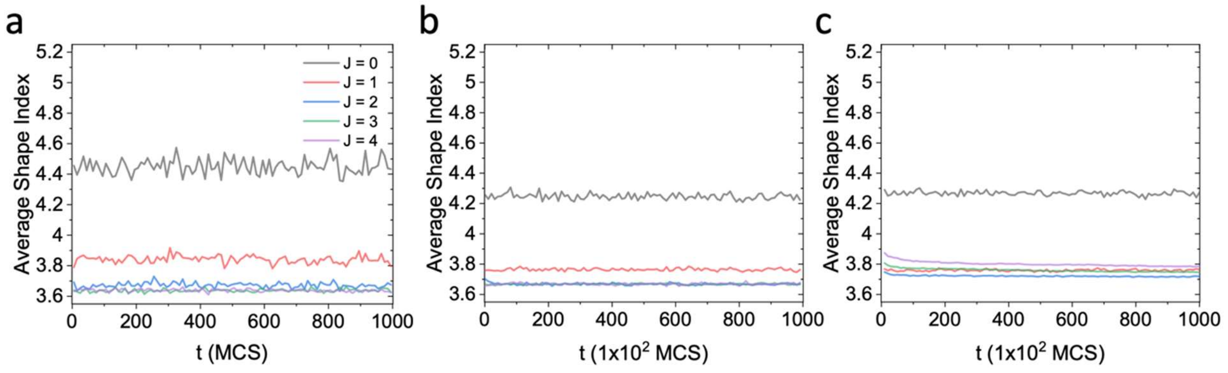


**Figure S4.** Average shape indices at each timestep in a representative simulation of (a) single cell, (b) aggregate, and (c) monolayer CPM simulations. The legend in (a) applies to all panels, where J denotes the relevant J parameter for each simulation type, as described in the main text.

**Figure S5**

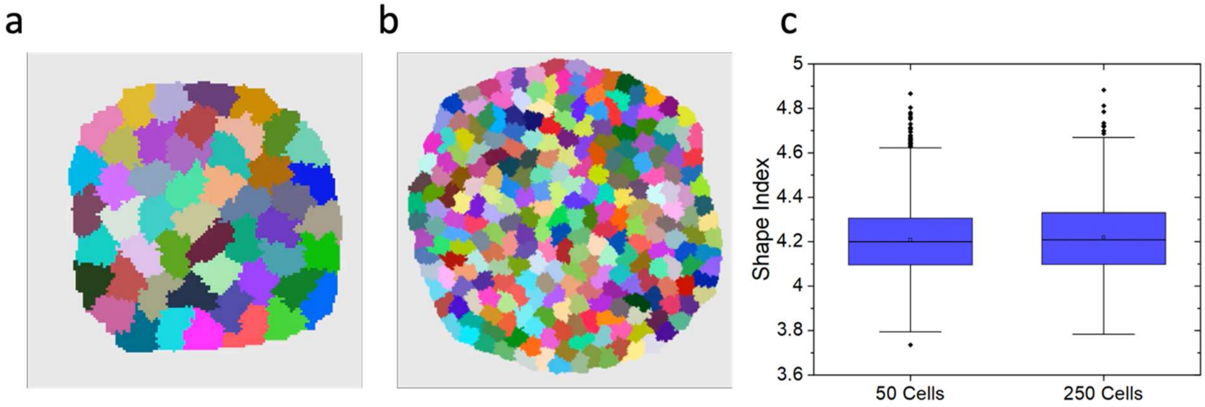


**Figure S5.** Cell shapes in CPM aggregates composed of (a) 50 or (b) 250 cells. (c) Shape distributions for each aggregate in (a-b) for $J_{cell} = 0$ and $J_{env} = 2$. To obtain similar numbers of measurements of cell shapes between differently sized aggregates, cells were pooled and measured over the final 10 MCS and 50 MCS, for 50 and 250 cell aggregates respectively (2500 cells each). As shown in Fig. S4, cell shapes do not vary significantly over time and can thus be pooled in this way without biasing measurements.

### Supporting Videos

#### Video 1

CPM simulations for representative single cells (top row, $1x10^3$ MCS total), aggregates (middle row, $1x10^5$ MCS total) and monolayers (bottom row, $1x10^5$ MCS total) for the same range of J values explored in Fig. 1 in the main text. For isolated cells, $J_{env}$ is varied from 0 to 3 (left to right) while for aggregates and monolayers $J_{cell}$ is varied over the same range (left to right). For aggregates, $J_{env} = 2$ as described in the main text.

**Video 2**

CPM simulations for an aggregate ($1x10^5$ MCS total length) with $J_{cell}$ = 1.75 and $J_{env}$ = 2.

**Video 3**

CPM simulations of round cell monolayers ($1x10^5$ MCS total length) over a select $J_{cell}$ range in the vicinity of the jamming transition. The transition from single cell motion to collective motion to jamming can be observed over this range by examining cell motion in the (top) full monolayer where different cells are represented by different colors and (bottom) by examining trajectories of a select group of cells (labeled in red) in the same monolayer.

**Video 4**

CPM simulations of (right) round and (left) elongated (left) cell monolayers ($1x10^5$ MCS total length) for $J_{cell} = 2$. Round cells are jammed while elongated cells display individual cell migration at this $J_{cell}$ value. (top) The full monolayers are shown and (bottom) select cells are highlighted to aid visualization.

**Video 5**

CPM simulations of elongated cell monolayers ($1x10^5$ MCS total length) for (left to right) $J_{cell}$ = 1.75, 2.00, 2.25, 2.50, and 2.75. Select cells are highlighted in red to highlight cell motion.
